# Slow-Timescale Regulation of Dopamine Release and Mating Drive Over Days

**DOI:** 10.1101/2025.05.29.656898

**Authors:** Praneel S. Sunkavalli, Joseph C. Madara, Lauren F. Christenson, Adriana Diaz, Janelle Chen, Luca Ravotto, Tommaso Patriarchi, Mark L. Andermann, Stephen X. Zhang

**Affiliations:** Division of Endocrinology, Diabetes and Metabolism, Beth Israel Deaconess Medical Center, Harvard Medical School, Boston, MA, USA; Institute of Pharmacology and Toxicology, University of Zurich, Zurich, Switzerland; Neuroscience Center Zurich, University and ETH Zurich, Switzerland; Center for Neural Science, New York University, New York, NY, USA

## Abstract

The rise and fall of motivational states may take place over timescales as long as many days. We used mouse mating behavior to model how the brain orchestrates slow-timescale changes in motivation. Male mice become sexually satiated after successful matings, and their motivation to mate gradually recovers over a week. Using deep-brain fluorescence-lifetime imaging in the medial preoptic area (MPOA), we found that tonic dopamine transmission—which regulates mating drive—also declined after mating and re-emerged over a week. Two mechanisms regulated dopamine transmission. First, successful mating transiently reduced tonic firing of hypothalamic dopamine-releasing neurons, thereby inhibiting dopamine release and mating behavior. Second, mating reduced the ability of these neurons to produce and release dopamine, and this ability gradually returned over the week-long recovery time course. Therefore, fast and slow mechanisms of neuronal plasticity cooperate to control the early and late phases of motivational dynamics, respectively.

## Introduction

The propensity and vigor of behaviors scale with internal states that reflect past experiences and competing options (Wei et al., 2021; Zhang et al., 2016, Zhang et al., 2021). These internal states can last seconds to days. The principles by which the brain controls behaviors over slower timescales are particularly poorly understood. One challenge to studying slow-timescale changes is the ability to quantify, in absolute terms, the slow changes in neuromodulator transmission that shape activity in behavioral circuits across internal states.

We used the mating behavior of male mice as a model to understand the mechanisms that govern slow-timescale changes in sexual motivation over days. After one or more successful matings, male mice have a greatly reduced interest in mating with receptive females, thereby instantiating a state of sexual satiety (Bayless et al., 2023; Zhang et al., 2021; Zhou et al., 2023), and this mating drive gradually recovers over the course of several days (McGill & Blight, 1963; Valente et al., 2021; Zhang et al., 2021; Zhou et al., 2023). Previously, we and others showed that dopamine transmission in the MPOA from nearby anteroventral and preoptic periventricular nuclei (AVPV/PVpo) is critical for controlling mating drive (Hull et al., 1986; Zhang et al., 2021). Here, we applied an analytic framework using fluorescence lifetime imaging microscopy (FLIM) and an existing neuromodulator sensor to quantify extracellular dopamine concentration in awake mice across days. We found that dopamine concentration in the MPOA is scaled down and gradually scaled back up across days in a manner that tracks the slow changes in mating behaviors. We identified two mechanisms – reduction and recovery in excitability of dopaminergic neurons and in dopamine synthesis – that jointly act as a multi-day timer for post-satiety recovery of mating drive via control of tonic dopamine release rate into the MPOA.

## Results

### Dopamine tone in the MPOA tracks recovery time course after sexual satiation

Sexual satiation following successful mating involves an evolutionarily conserved state of reduced mating drive (Bayless et al., 2023; Beach & Jordan, 1956; McGill & Blight, 1963; Zhang et al., 2016, Zhang et al., 2021; Zhou et al., 2023) (Figure 1A). In home-cage mating experiments using male C57BL/6 mice, we found that this switch took place after one or two matings, as observed by suppression of both sniffing and mounting behaviors (Figures 1B-1C) (Bayless et al., 2023; Zhang et al., 2021; Zhou et al., 2023). Once satiated, the motivation to mate was not re-invigorated by pairing with new females. Instead, male mice gradually recovered from the satiated state over ∼7 days of abstinence (Figures 1B-1C). The time course of behavioral recovery qualitatively tracks changes in sperm counts following satiation, which partially recovered after 7 days (Figure S1A). In contrast, the social behaviors associated with intruder identification were not affected (Figure 1D). These results suggest the possibility that male mice calibrate their mating behavior to reproductive potency over days of recovery (Zhang et al., 2016).

**Figure 1.**
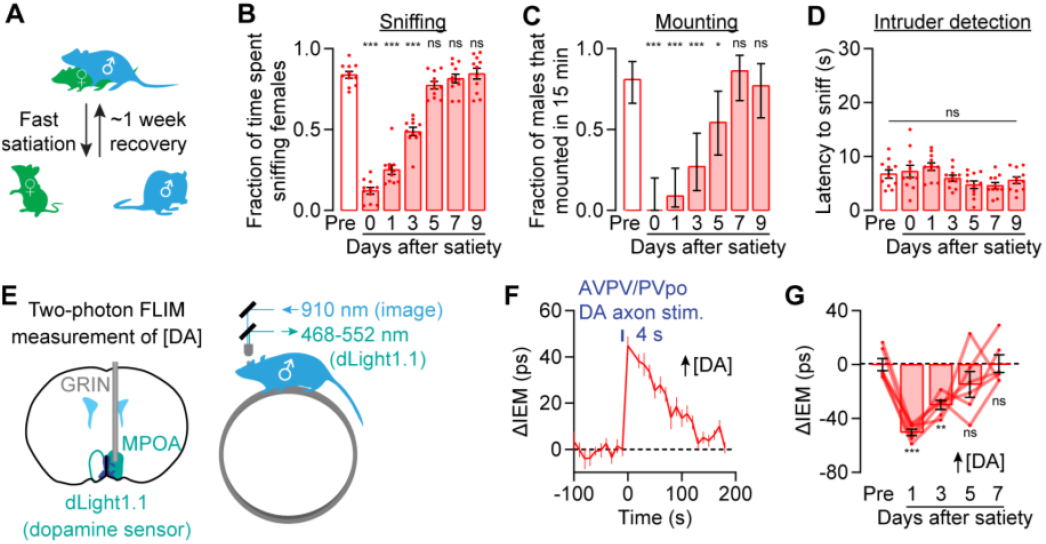
Recovery of male mating drive across days. **(A-C)** After sexual satiation, both the fraction of time that males engaged in sniffing behavior during the pre-mounting period (first 5 minutes after female entry) (B) and the fraction of males that attempted to mount the female within 15 minutes after entry (C: mean ± 95% c.i.) decreased and gradually recovered within ∼5-7 days. N = 11 male mice. See also Zhang et al., 2021. **(D)** The latency to sniff an intruder placed in the home cage was not changed after sexual satiation. N = 11 male mice. **(E)** *In vivo* 2p-FLIM imaging of tonic dopamine levels in MPOA using the sensor dLight 1.1 via a GRIN lens. **(F)** AVPV/PVpo dopamine axon stimulation in the MPOA of a baseline (non-satiated) mouse induces an elevation in dopamine as determined using the integral extraction method (IEM) to estimate dLight1.1 fluorescence lifetimes. Dopamine levels gradually return to baseline over minutes. N = 9 Fields of View (FOVs) from 3 males. **(G)** IEM quantification of dLight1.1 fluorescence lifetime in the MPOA showed a decrease from the pre-satiety quantification (calculated by averaging dLight1.1 fluorescence lifetime of 3 pre-satiated days) in the absolute dopamine concentration in this brain region that gradually recovered within ∼1 week. N = 6 FOVs from 3 males. Mean ± s.e.m. *p<0.05, **p<0.01, ***p<0.001. See Supplementary Table 1 for statistics.

Dopamine release in MPOA from nearby AVPV/PVpo nuclei is critical for controlling mating drive in male mice (Zhang et al., 2021). Previous experiments examining the expression of the immediate early gene *c-fos* suggest that these dopamine neurons are tonically active when males have high mating drive and are quiescent in the satiated state (Zhang et al., 2021). Satiation also curbs the phasic dopamine transients in the male mouse’s MPOA that occurs when the animal approaches a female (Zhang et al., 2021).

Systemic or MPOA-specific infusion of dopamine agonists is sufficient to induce mating behaviors in otherwise sexually satiated male animals (Mas et al., 1995; Rodríguez-Manzo, 1999). We therefore investigated the possibility that satiation and recovery of mating behaviors are timed by the reduction and subsequent replenishment of tonic dopamine transmission in the MPOA.

To measure absolute dopamine concentrations in the MPOA of awake male mice, we used two-photon fluorescence lifetime imaging microscopy (2p-FLIM) to image the dopamine sensor dLight1.1 through an implanted GRIN lens (Zhang et al., 2021, Zhang et al., 2024). We chose this technique because 1) lifetime measurements allowed us to estimate absolute sensor states, 2) an imaging-based technique allows for sampling of dopamine concentrations throughout a 3D volume rather than at a single point, and 3) the spatial resolution of two-photon imaging allowed us to reduce contamination by autofluorescence (see Methods) (P. Ma, Sternson, et al., 2024). We considered two potential methods of lifetime quantifications: the average arrival time (Tm) and the integral extraction method (IEM) (Figure S2A and see Method for details). Tm is a commonly used metric with a superior signal-to-noise ratio (SNR) (P. Ma, Sternson, et al., 2024; Zhang et al., 2021, Zhang et al., 2024). However, Tm is also influenced by fluorescence intensity changes such that, for common sensors whose bound-state is brighter than the apostate, the dynamic range of the optimal ligand concentrations is left-shifted (i.e., left-shifted apparent Kd) (Figures S2B-S2D). Therefore, for intensity-optimized sensors (e.g. GCaMP), Tm is prone to saturation and shows suboptimal lifetime changes (P. Ma, Chen, et al., 2024). IEM is influenced by fluorescence intensities in a manner that is mathematically distinct from Tm and reports a less left-shifted Kd (Figure S2E). However, IEM has a lower SNR than Tm (Figure S2F-S2I) and is therefore better suited for settings where each lifetime estimate is derived using a high photon count.

We compared these two metrics by using 2p-FLIM to measure dopamine transients that are evoked by optogenetic stimulation of dopamine axons from AVPV/PVpo nuclei (Figure S3A). By pooling the photons in each field of view, we calculated Tm and IEM for dLight1.1 using 300,000-1,000,000 photons per lifetime. We found that, under these conditions, the IEM metric consistently produced greater lifetime changes than Tm for dLight1.1, presumably because the range of dopamine concentrations in this region are more closely aligned with the dynamic range of IEM than Tm quantification (Figures 1E-1F and S3B-S3D).

Following this validation, we used 2p-FLIM to quantify dopamine concentration at different imaging depths in the MPOA across days following sexual satiation (Figure S3E-S3G). Both IEM and Tm quantification methods indicated a significant reduction in dopamine concentration in the first two days following satiation (Figures 1G and S3E-S3F), followed by a gradual recovery to baseline within about a week. These observations could not be made using sensor intensity quantification, which is more sensitive to day-to-day variations in imaging conditions (Figure S3G). These results demonstrate parallel slow-timescale changes in tonic dopamine transmission and in mouse behavior after sexual satiety, supporting the hypothesis that changes in tonic dopamine transmission may contribute to behavioral recovery over days.

### Differential changes in spike rates and dopamine synthesis during recovery

One potential cause for changes in tonic dopamine transmission is the change in tonic spike rates of the dopaminergic neurons, a hypothesis that is supported by previous findings of mating state-dependent *c-fos* expression (Zhang et al., 2021). To directly record spike rates, we performed cell-attached recordings of AVPV/PVpo neurons in brain slices from sexually satiated mice (Figure 2A). Most AVPV/PVpo dopamine neurons fired spontaneous action potentials in brain slices, and their firing rates were significantly decreased after satiation (Figure 2B). However, by Day 2 of recovery, when mice were still uninterested in mating, the firing rates of these dopamine neurons had already recovered to baseline (Figure 2C). The decrease and recovery of firing rates were seen in dopamine neurons sampled across AVPV and PVpo (Figures 2D-2F) and were not different between the two areas (Figures 2G-2H). Therefore, restoration of the dopamine neurons’ spontaneous firing rates took place much faster than the observed behavioral 7-day recovery in mating drive (Figure 2I).

**Figure 2.**
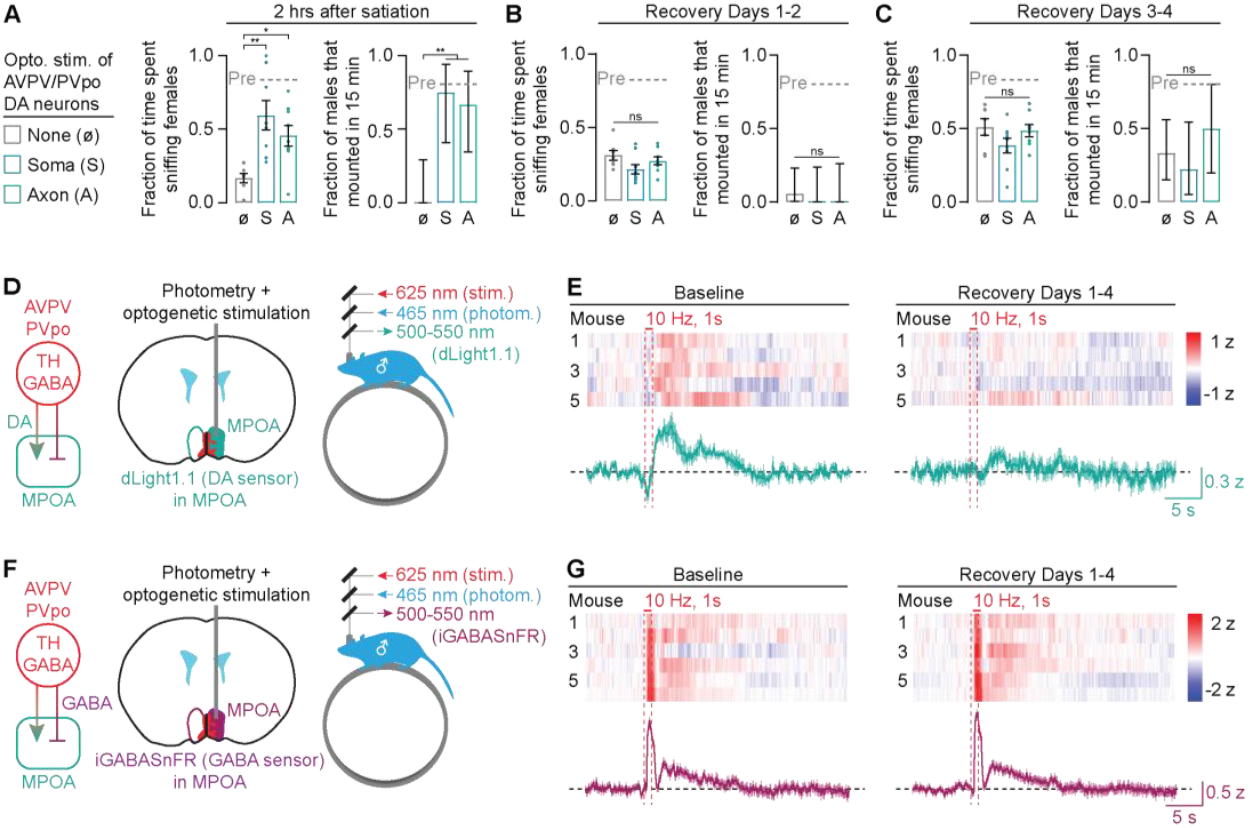
Fast drop and recovery of dopaminergic AVPV/PVpo neuron firing after sexual satiation. **(A-C)** In cell-attached patchclamp recordings of dopaminergic AVPV/PVpo neurons, spontaneous firing rates decreased overnight after satiation (A, B) but recovered by Day 2 (C). A Cre-dependent mCherry was injected bilaterally to identify dopaminergic AVPV/PVpo neurons. **(D-F)** Firing rates of AVPV and PVpo dopamine neurons recorded in male mice prior to satiation (D: N = 43 neurons from 7 males), 1 day post satiation (E: N = 58 neurons from 8 males), and 2 days post satiation (F: N = 48 neurons from 7 males). Firing rates (scaled by disc size) of neurons are mapped onto their coronal locations. **(G-H)** The changes in firing rate of dopaminergic neurons in AVPV (G: N = 28 neurons from 7 males [Baseline], 36 neurons from 8 males [Recovery Day 1], 15 neurons from 7 males [Recovery Day 2]) and PVpo (H: N = 15 neurons from 7 males [Baseline], 22 neurons from 8 males [Recovery Day 1], 33 neurons from 7 males [Recovery Day 2]). **(I)** The recovery of dopaminergic AVPV/PVpo neuron firing rates does not parallel the return of sexual behaviors and absolute dopamine concentration in the MPOA. Mean ± s.e.m. *p<0.05, **p<0.01, ***p<0.001. See Supplementary Table 1 for statistics.

To test the role of AVPV/PVpo dopamine neurons in controlling the recovery time course, we optogenetically stimulated their somas and their axons in the MPOA at different time points after satiation. Within two hours after satiation, both axonal and somatic photostimulation (10 Hz, 1 mW) were sufficient to acutely re-invigorate the sniffing and mounting behaviors of males paired with new females (Figure 3A). However, in experiments throughout the subsequent four days, the same stimulation protocol could no longer elevate mating behaviors beyond the pace of natural recovery (Figure 3B-3C). Together with the electrophysiological recordings above, these behavioral experiments suggest an hours-long, post-sa-tiation window in which mating drive could be re-invigorated by elevating the firing rate of the dopamine neurons. Beyond this point, however, an additional layer of regulation titrates the gradual recovery in mating drive.

**Figure 3.**
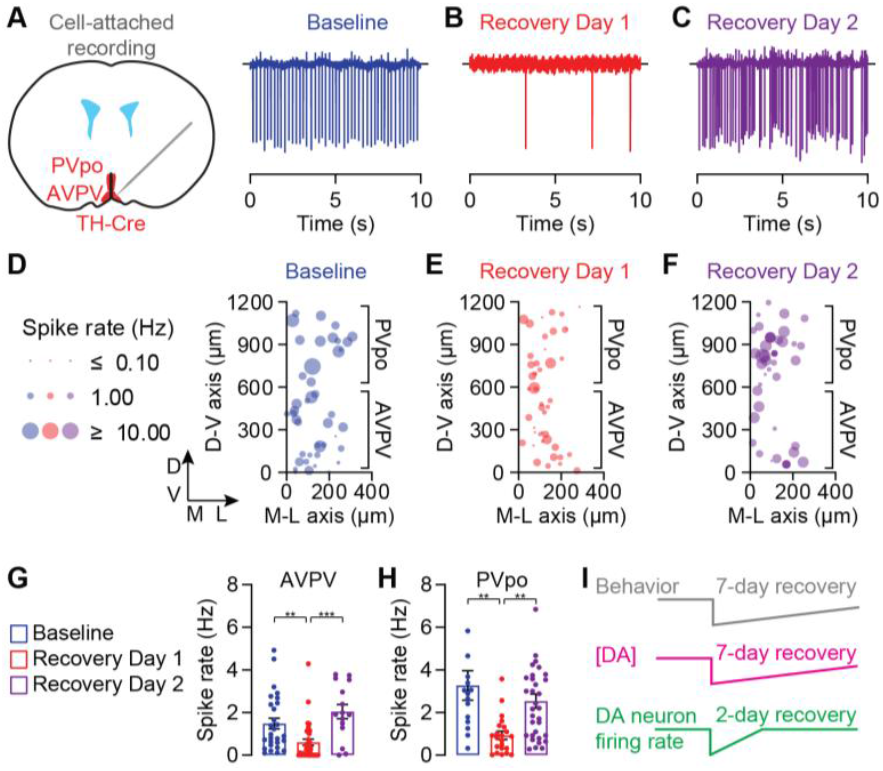
Selective reduction of transmission of dopamine but not GABA from AVPV/PVpo dopaminergic neurons after sexual satiation. **(A-C)** Optogenetic stimulation of dopaminergic somas in AVPV/PVpo or their axons in MPOA increased sniffing and mounting behaviors two hours after sexual satiation (A) but not at later times in the subsequent four days (B, C). For comparison, pre-satiety measurements are indicated by grey dashed lines (see Figure 1). The LED pulses were turned on 1 min before female entry and turned off 5 min after female entry. Data in 3A were replotted from Zhang et al. N = 7-9 male mice. Left panels of A-C: mean ± s.e.m.; right panels: mean ± 95% c.i. **(D-E)** Brief *in vivo* photostimulation of dopaminergic AVPV/PVpo axons in the MPOA (D) triggered a dopamine transient (measured by dLight1.1 photometry) that is weakened in the first few days following sexual satiation (E). Heatmaps show mean traces of 3 baseline days and 4 post-satiation days for each mouse. Baseline data were replotted from Zhang et al., 2021. N = 5 mice. **(F-G)** Brief *in vivo* photostimulation of dopaminergic AVPV/PVpo axons in the MPOA (F) drives GABA release (measured by iGABASnFR photometry) that remains unchanged following sexual satiation (G). Heatmaps show mean traces of 3 baseline days and 4 post-satiation days for each mouse. N = 6 mice. Mean ± s.e.m. *p<0.05, **p<0.01, ***p<0.001. See Supplementary Table 1 for statistics.

Having ruled out a recovery timer that is based solely on firing rates of AVPV/PVpo dopamine neurons, we next investigated the ability of these neurons to release dopamine. In the hypothalamus, most dopamine neurons are developmentally related to GABA neurons and, in adults, most of these dopamine neurons also express the GABA vesicular transporter VGAT (Negishi et al., 2020; Romanov et al., 2020). We therefore tested the possibility that the AVPV/PVpo dopamine neurons co-release dopamine and GABA.

We used fiber-photometry and the dopamine sensor dLight1.1 to record dopamine release in the MPOA that is evoked by photostimulating AVPV/PVpo dopamine axons (Figure 3D). In the baseline state, a 1-s photostimulation train evoked a dopamine transient that lasted ∼20 seconds (Figure 3E and S4A). However, in the first few days after satiation and in the same mice, the same stimulation protocol evoked much weaker transients, indicating a reduction in dopamine transmission. When we used a GABA sensor (iGABASnFR) (Marvin et al., 2019) to measure GABA release from the same axons, we observed a biphasic response in the MPOA: an early peak lasting ∼1 second followed by a late peak lasting ∼20 seconds (Figure 3F-3G). Because the two peaks were correlated on a trial-by-trial basis (Figure S4B-S4C) and ∼81% of MPOA neurons are GABAergic (see Moffitt et al., 2018), we interpreted the first peak as the initial GABA release from the optogenetic axon stimulation, and we speculate that the second peak reflects the GABA release from the dopamine-driven activation and/or rebound excitation of local MPOA neurons. After sexual satiation, at a time point when dopamine transmission was reduced, the sizes of both iGABASnFR peaks remained largely unchanged (Figure 3G and S4D-S4E), indicating intact GABA transmission and axonal excitability after satiation. These results argue that sexual satiety induced a selective reduction in dopamine (but not GABA) transmission from AVPV/PVpo axons in the MPOA that lasts several days.

### Dynamic regulation of neurotransmitter composition as a multi-day timer

Given that dopamine but not GABA transmission is reduced after sexual satiety, we looked for dopamine-specific mechanisms underlying this change. In female mice (but not in males), tyrosine hydroxylase (TH) – the * rate-limiting enzyme in dopamine biosynthesis – is expressed in more AVPV neurons following the onset of parenthood (Scott et al., 2015). We hypothesized that TH expression in AVPV neurons can be reduced and reinstated over the course of male sexual satiation and recovery. Because AVPV and PVpo are small nuclei containing both dopaminergic and other neurons, we decided to quantify TH expression after clearing intact tissue using iDisco (Renier et al., 2014) (Figures 4A and S5A) and staining for TH. We scanned 3D volumes of tissue using two-photon microscopy and segmented the TH-expressing somas using Cellpose 2.0 (Supplementary Video 1). In the AVPV, TH expression remained dense in the days following satiation, and no significant changes in the number of TH+ cells were observed (Figure 4B). In the PVpo, the number of TH+ cells was reduced by 72% in the first 3 days after sexual satiety, and its expression gradually re-emerged by 7 days post-satiation (Figure 4B). These dynamics match the recovery time course of extracellular dopamine concentration in the MPOA and the recovery of males’ mating behaviors (Figure 1).

**Figure 4.**
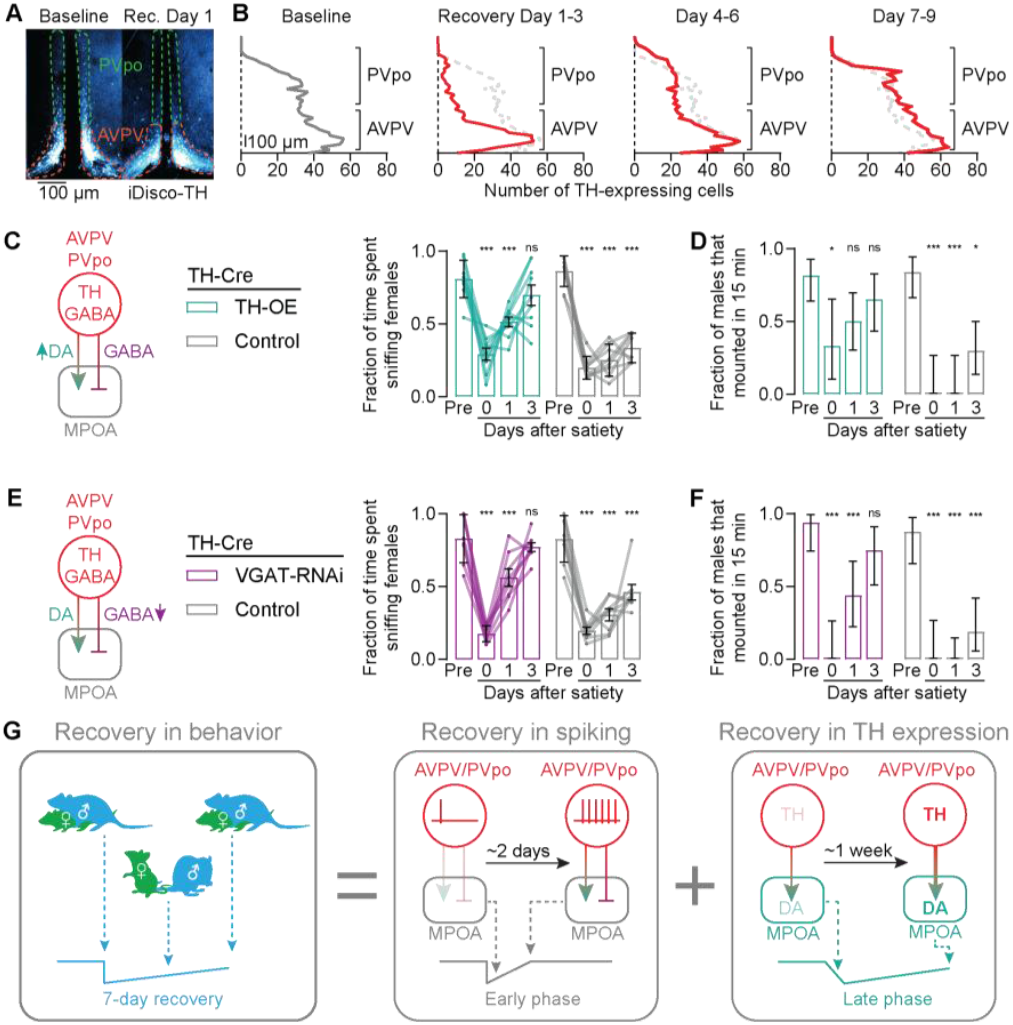
Tyrosine Hydroxylase (TH) expression controls the multi-day recovery of mating drive. **(A)** Quantification of TH protein expression in the somas of AVPV/PVpo neurons using iDISCO. Red rectangle denotes the location of PVpo where TH expression appeared satiety-dependent. **(B)** After sexual satiation, TH expression decreased and gradually recovered in PVpo over the 7-9 days of recovery. For comparison, pre-satiety cell counts are replotted as dashed lines in the subsequent panels. N = 10 male mice [Baseline], 9 male mice [Recovery day 1-3], 7 male mice [Recovery day 4-6], 7 male mice [Recovery day 7-9]. **(C-D)** Viral overexpression of TH (AAV-EF1a-DIO-TH-p2A-mRFP) in AVPV/PVpo dopamine neurons accelerated the recovery of sniffing (C) and mounting behaviors (D) after satiation. C: mean ± s.e.m.; D: mean ± 95% c.i. N = 9 male mice. **(E-F)** RNAi knockdown of VGAT (AAV-hSyn-FLEX-dsRed-shVgat) in AVPV/PVpo dopamine neurons accelerated the recovery of sniffing (E) and mounting behaviors (F) after satiation. E: mean ± s.e.m.; F: mean ± 95% c.i.; N = 8 male mice. **(G)** A dual-component timer underlies the recovery of sexual motivation. First, the spike rate of AVPV/PVpo dopamine neurons is reduced after satiation, and it recovers after ∼2 days. Second, the expression of TH decreases after satiation, and it recovers gradually over ∼1 week. Mean ± s.e.m. p<0.05, **p<0.01, ***p<0.001. See Supplementary Table 1 for statistics.

By tuning TH expression, a dopaminergic neuron may be able to flexibly alter its neurotransmitter composition and change the ratio of dopamine-to-GABA transmission. We therefore tested the impact of artificially manipulating the ratio of dopamine/GABA transmission to MPOA on mating-drive recovery. We first used a previously validated AAV that overexpresses TH in Cre-expressing neurons (TH-OE) (Scott et al., 2015). Because this construct uses the exogenous EF1α promoter, it is presumably not regulated by the same transcriptional factors that control endogenous TH expression. In the baseline state, TH-OE mice displayed sniffing and mounting behaviors that are comparable to control mice (Figures 4C-4D). However, these mice recovered significantly faster from sexual satiety, reaching nearbaseline levels of mating behaviors in ∼3 days (Figures 4C-4D). These results suggest that artificial TH overexpression may hasten this behavioral recovery by dampening the behavioral sensitivity to the endogenous fall in TH expression and dopamine transmission.

To test the impact of GABA co-transmission, we used a different AAV that uses a Cre-dependent short-hairpin RNA to knock down the expression of the vesicular GABA transporter VGAT by ∼60% (VGAT-RNAi) (Yu et al., 2015). In AVPV/PVpo axons, this construct should reduce GABA transmission by limiting its packaging rate into synaptic vesicles, thereby increasing the ratio of dopamine-to-GABA release into the MPOA. Similar to TH-OE, knocking down VGAT also hastened the recovery time course in a manner that was not seen in mice that expressed the scrambled control (Figures 4E-4F). These results show that artificially increasing dopamine-to-GABA ratio by either increasing dopamine production or decreasing GABA packaging can hasten the time course of recovery of mating drive.

## Discussion

Taken together, our results argue for a two-component timer that regulates the gradual changes in dopamine transmission in the MPOA and in mating behavior over the 7 days of mating-drive recovery. First, the tonic firing rates of AVPV/PVpo dopamine neurons are decreased after sexual satiation to acutely reduce dopamine release in the MPOA, and this state of inactivity lasts for 1-2 days (Figure 2). Second, the capacity of AVPV/PVpo dopamine neurons to synthesize and release dopamine is reduced by diminishing the expression of TH, which then gradually returns to baseline over ∼7 days (Figures 3-4). Because the tonic firing rates of these neurons can be decreased much more rapidly than the removal of TH proteins (Tekin et al., 2014), this drop in excitability is likely responsible for mediating the initial onset of sexual satiety after successful mating, similar to previous chemogenetic inhibition experiments using hM4Di (Zhang et al., 2021). In this early phase and before TH removal, optogenetic stimulation of AVPV/PVpo dopamine neurons can revert sexual satiety (Figure 3A-3C). Once TH levels diminish, stimulation of AVPV/PVpo dopamine neuron cell bodies (or axons in the MPOA) can no longer evoke dopamine release or revert the state of satiety (Figure 3A-3E). At this late recovery stage, it is the time course of re-expression of TH (Figure 4A-4B), rather than the time course of restoration of tonic firing rates (Figure 2), that matches the recovery of dopamine transmission and mating behaviors (Figure 1). These findings highlight that changes in the spiking activity of AVPV/PVpo dopamine neurons alone only partially capture behavioral dynamics over days. Therefore, these two mechanisms may be complementary in mediating the early and late phases of post-mating satiety (Figure 4G).

Neurotransmitter switching by altering TH expression has been observed in different behavioral contexts (Aumann et al., 2013; Dulcis et al., 2013, 2017; Frank et al., 2023; Scott et al., 2015; Spitzer, 2015). In the AVPV/PVpo dopamine neurons, sexual satiation weakened dopamine transmission, but not GABA transmission, so the term neurotransmitter switching may only apply partially. Nevertheless, we believe the plasticity mechanisms that regulate TH expression are the same as those driving neurotransmitter switching in developing and adult animals (Spitzer, 2017). In adult animals, activity-dependent TH transcription is thought to be the driver of opposite-direction shifts in the expression of TH and somatostatin in the rodent paraventricular and periventricular hypothalamus during altered photoperiods, as well as changes in TH-expression in the *Xenopus* spinal cord (Dulcis et al., 2013; Pierani et al., 2001). We currently do not know which step(s) in TH transcription and/or translation are regulated during recovery in the AVPV/PVpo dopamine neurons. It is possible, that the drop and subsequent rise in tonic firing rates of AVPV/PVpo dopamine neurons actually promote the observed changes in TH expression. In this way, excitability and molecular changes in the AVPV/PVpo dopamine neurons may be controlled by the same synaptic inputs. However, since the reexpression of TH took place over ∼5 days, it is unclear whether activity-dependent transcription alone could be responsible for such a slow change, and other modes of event-triggered gene regulation may be involved (Abraham et al., 2019).

In contrast to dopamine, GABA co-transmission from the AVPV/PVpo dopamine neurons is unaffected by sexual satiety (Figures 3F-3G). Functionally, GABA co-transmission from these neurons suppresses mating behavior (Figures 4E-4F) and thus opposes the actions of dopamine (see also Tritsch et al., 2016). Therefore, in this push-pull system, we believe that the role of GABA co-transmission is to set a threshold that determines the degree of dopamine release needed for a net mating-promoting effect (Zhang et al., 2021). This threshold may underlie the observation that recovery of tonic firing without full restoration of TH expression is insufficient to promote the recovery of mating drive (i.e., on Day 2 of recovery). The nature of this thresholding effect is, however, not currently understood. This is because we do not know the receptor type through which GABA actions in MPOA suppress mating drive, and their possible interaction with cAMP-mediated signaling downstream of D1-family dopamine receptors in the MPOA (Zhang et al., 2021). We also do not know whether these two co-transmitters are in fact received by the same downstream neurons, and how the expression of their receptors may be maintained and altered during recovery. Further research is required to understand receptor expression patterns, signal integration mechanisms, and possible post-satiety plasticity at the level of receptor expression.

Outside of AVPV/PVpo dopamine neurons, there are other cells and mechanisms that are also important for sexual satiety. Recently, Zhou and colleagues showed that Esr2-expressing neurons in the BNST become persistently hyper-active during sexual satiety and suppress mating (Zhou et al., 2023). The hyperexcitability is due to elevated HCN currents that are different between baseline, satiated, and recovered states. Another study demonstrated that stimulating Tac1-expressing neurons in the BNST (BNST^TAC1^) or TAC1 receptor-expressing neurons in the POA can temporarily reinvigorate sexual behaviors post-satiety (Bayless et al., 2023). Since these BNST^TAC1^ neurons are not dopaminergic, important insights may be gained from day-by-day investigations of potential timing mechanisms that arise from their parallel regulation of POA, possibly also through Gs-coupled GPCR signaling. Moreover, whereas chemogenetic inhibition of the AVPV/PVpo dopamine neurons significantly reduces mating drive (Zhang et al., 2021), partial ablation of AVPV/PVpo dopamine neurons (∼50%-67% neuron loss) and their fibers innervating the MPOA only transiently reduces mating behaviors for ∼1 day (Bayless et al., 2023; Bazzett et al., 1992; Scott et al., 2015). After that, the circuit is functional but more sensitive to other methods of dopamine-synthesis disruption (Bazzett et al., 1992). Together, these studies suggest that sexual satiety likely recruits plasticity mechanisms in multiple mating-related circuits, the effect of which can be overcome by experimentally activating key circuit nodes.

*In vivo* fluorescence lifetime measurements using 2p-FLIM and one-photon fluorescence lifetime imaging photometry (1p-FLIP) are emerging as a powerful new frontier in technological development that enables longitudinal quantification of biochemical and biophysical parameters in the brain across hours and days (Díaz-García et al., 2017; Gautham et al., 2024; Lee et al., 2019, 2021; Lodder et al., 2025; L. Ma et al., 2022; P. Ma, Chen, et al., 2024; Zhang et al., 2021, Zhang et al., 2024). Three important areas for further improvement are instrumentation, sensor development, and analysis pipelines. In this work, we applied an expanded framework for *in vivo* fluorescence lifetime analysis and demonstrated its practical utility in enabling the lifetime recording of dLight1.1 (P. Ma, Chen, et al., 2024). This sensor is neither new nor optimized for FLIM, but the analysis approach employed in this study should also be applicable to the rapidly emerging library of FLIM-optimized tools (see also Lodder et al., 2025). We demonstrated that Tm is a high-SNR method of lifetime quantification that nevertheless left-shifts the apparent Kd of the sensor, while IEM can be applied to estimate lifetimes with less Kd shifting in situations with high photon counts. We envision that the choice between Tm and IEM methods will be made on an experiment-by-experiment basis, as FLIM sensors and hardware continue to improve. Future development in autofluorescence rejection algorithms will also be critical for translating fluorescence lifetime estimates to biological units (e.g., μM) (P. Ma, Sternson, et al., 2024). Our study highlights how new fluorescence lifetime techniques can help reveal mechanisms underlying changes in behavior across the rarely explored timescale of days.

## Acknowledgements

We thank B. Lowell, M. Crickmore, D. Rogulja, J. Blau, A. Nelson, D. Lin, A. Falkner, J.S. Alvarado, X. Cai, P. Kalugin, M. Porniece, J. Stern, B. Lodder, T. Kamath, and members of the Zhang lab for useful feedback. M. Hammell, J. Baker, S. Sankar, J. Chen, P. Prasad, D. Guarino, H. Choh, A. Sambangi, H. Lauterwasser, C. McHugh, D. Fleharty, H. Kaul, and K. Fernando for helping with animal care, behavioral experiments and histology. Boston Children’s Hospital Viral Core provided viral services. O. Yizhar provided us with the TH-OE AAV vector. Authors were supported by an HMS Broderick Phytocannabinoid Research Grant, Simons Foundation Pilot Award ID 976096, and two Harvard Brain Science Initiative Bipolar Disorder Seed Grants, and K. and L. Dauten (M.L.A.); Lefler Fellowship, Charles A. King Trust Fellowship, and NIH K99 DK134853 (S.X.Z.). Figure 3A-3C and Figure 3D-3E were reproduced from data collected in Zhang et al. 2021.

## Author contributions

S.X.Z. conceived the project. P.S.S. and S.X.Z. wrote the manuscript with methodological and editing inputs from M.L.A. and all other authors. P.S.S. and S.X.Z. conducted all surgeries. P.S.S., L.F.C., A.D., J.C., and S.X.Z. performed the behavioral experiments and fiber photometry recordings. P.S.S., L.F.C., and S.X.Z. performed the in vivo two-photon FLIM experiments. J.C.M performed electrophysiology experiments. P.S.S., L.F.C., A.D., and J.C. performed all histology. All authors contributed to data analysis.

## Materials and Methods

### Animals

All animal care and experimental procedures were approved by the Institutional Animal Care and Use committee at Beth Israel Deaconess Medical Center (BIDMC). Animals were house in a 12-hour light/12-hour darkness environment (temperature 20-24 °C, humidity 30-70%) with standard mouse chow and water provided ad libitum, unless specified otherwise. Male and female mice older than 8 weeks were used in experiments, but due to the scope of the study, recordings, perturbations, and behavioral analyses were focused on males. We used the following genotypes: male and female C57BL/6J (WT; 000664, The Jackson Laboratory), male B6.Cg-7630403G23RikTg(Th-cre)1Tmd/J (TH-Cre; 008601, The Jackson Laboratory) (Black et al., 1998; Champy et al., 2008; Drake et al., 2001; Fontaine & Davis, 2016; Ishida et al., 1991; Kapur et al., 2020; Mouse Genome Sequencing Consortium, 2002, p. 129; Paigen et al., 1985; Petkov et al., 2004; Savitt et al., 2005; Simon et al., 2013; Toye et al., 2005; White et al., 2002), and their F1 progeny. We used TH-Cre mice to label AVPV/PVpo dopamine neurons because these cells only sparsely express DAT (Slc6a3) (Lein et al., 2007) and cannot be targeted using DAT-Cre mice. Males’ mating behaviors and ability to process sensory cues from conspecifics are strongly improved by having at least some social experience (Y. Li et al., 2017; Remedios et al., 2017; Swaney et al., 2012). Hence, we paired males individually with hormonally primed females, two weeks before any experiment. This two-week time frame was chosen to be longer than recovery time from sexual satiety (Figures 1A-C). Once the female was removed, the male was single-housed in his home cage and remained so throughout the experiments. Mice with implants (e.g., fiber, GRIN lens) were single-housed to avoid damage. Sample sizes were chosen to reliably measure experimental parameters and keep with standards of the relevant fields (Zhang et al., 2021), while remaining in compliance with the ethical guidelines to minimize the number of experimental animals. Experiments did not involve experimenter-blinding.

### Stereotaxic surgeries

Viral injections, fiber implantations and GRIN lens implantations were generally performed as described in Zhang et al. (Zhang et al., 2021, Zhang et al., 2024) with the following specifications. For experiments that used a 400-μm diameter fiber or 500-μm diameter GRIN lens, we pre-set the insertion tracks by lowering needles with matching diameters (27 gauge and 25 gauge, respectively) to a depth of 0.1 mm above the final depth of the fiber or lens in order to ensure a snug fit for the fiber or lens, reduce brain motion, and accelerate recovery. All AAVs were injected at a titer of 3-15 1013 gc/ml (see below for volumes used). In the minority of cases where viral expression was absent (<10% of surgeries), we excluded the data from subsequent analyses. All animals were allowed to recover for at least 3 weeks prior to the onset of the experiments. No obvious capsid competition was seen in histology.

For optogenetic stimulation of AVPV/PVpo dopamine axons in the MPOA, AAV1-EF1-dfloxhChR2(H134R)-mCherry-WPRE-hGh (150 nl; AV-1-20297P, Penn Vector Core) was bilaterally injected into the AVPV of TH-Cre mice (Bregma: AP 0.25 mm, ML ±0.3 mm, DV −5.5 mm). An optic fiber with a metal ferrule (400 μm diameter core, multimode, 6.0 mm length, NA 0.48, Doric or homemade) was implanted unilaterally into the MPOA (Bregma: AP −0.16, ML 0.4 mm, DV −5.3 mm).

For optogenetic stimulation of AVPV/PVpo dopamine somas, AAV1-EF1-dflox-hChR2(H134R)-mCherry-WPRE-hGh was unilaterally injected into the AVPV of TH-Cre mice (150 nl; Bregma: AP 0.25 mm, ML 0.3 mm, DV −5.5 mm). An optic fiber (200 μm diameter core, multimode, 6.0 mm length, NA 0.48, Doric) was implanted ipsilaterally above PVpo (Bregma: AP 0.0 mm, ML 0.2 mm, DV −5 mm).

For dLight1.1 photometry experiments, AAV1-Syn-Flex-ChrimsonR-tdTomato (150 nl; UNC viral core) was injected bilaterally into the AVPV/PVpo (Bregma: AP 0.25 mm, ML ±0.3 mm, DV −5.5 mm) of TH-Cre mice. AAV2/1-Syn-dLight1.1 (Boston Children’s Hospital Vector Core) was injected unilaterally into the MPOA (150 nl; Bregma: AP −0.16 mm, ML 0.4 mm, DV −5.5 to −5.35 mm) and an optic fiber (400 μm diameter core, multimode, 6.0 mm length, NA 0.48, Doric) was implanted in the MPOA, ipsilateral to the dLight1.1 injection (Bregma: AP −0.16, ML 0.4 mm, DV −5.3 mm).

For iGABASnFR photometry experiments, AAV1-Syn-Flex-ChrimsonR-tdTomato (75 nl; Addgene) was injected bilaterally into the AVPV/PVpo (Bregma: AP 0.25 mm, ML ±0.3 mm, DV −5.5 mm) of TH-Cre mice. A 1:1 mixture of AAV1-hSyn-FLEX-iGABASnFR (Addgene 112163-AAV1) and AAV5-hsyn-Cre (Addgene 105553-AAV5) was injected unilaterally into the MPOA of TH-Cre mice (75 nl total; Bregma: AP −0.16 mm, ML 0.4 mm, DV − 5.5 to −5.35 mm). An optic fiber (400 μm diameter core, multimode, 6.0 mm length, NA 0.48, Doric) was implanted in the MPOA, ipsilateral to the iGABASnFR injection (Bregma: AP −0.16, ML 0.4 mm, DV −5.3 mm).

For cell-attached recordings of AVPV/PVpo neurons, AAV1-Syn-DIO-mCherry (150 nl; Addgene) was injected bilaterally into the AVPV/PVpo (Bregma: AP 0.25 mm, ML ±0.3 mm, DV −5.5 mm) of TH-Cre mice.

For experiments involving two-photon fluorescence lifetime imaging (FLIM) of dLight1.1 in the MPOA during recovery from sexual satiety in the head-fixed males, AAV1-syn-Flex-Chrimson-dTomato (150 nl; UNC Vector Core) was injected bilaterally into the AVPV/PVpo (Bregma: AP 0.25 mm, ML ±0.3 mm, DV −5.5 mm) of TH-Cre mice. AAV2/1-Syn-dLight1.1 (150 nl; Boston Children’s Hospital Vector Core) was injected unilaterally into the MPOA (Bregma: AP −0.16 mm, ML 0.4 mm, DV −5.5 to −5.35 mm). A doublet GRIN lens (NEM-050-25-10-860-DM, Grintech; 0.5 mm diameter; 9.89 mm length; 250 μm focal distance on brain side at 860 nm [NA 0.5]; 100 μm focal distance on the air side [NA 0.19]; 2.6x magnification through the lens) was implanted in the MPOA, ipsilateral to the site of expression of dLight1.1 (Bregma: AP −0.16, ML 0.5 mm, DV −5.3 mm). Given our previous behavioral data and intracellular-signal recording (Zhang et al., 2021), we interpret dLight1.1 signals in the MPOA as faithfully reporting dopamine release. However, since dLight1.1 fluorescence intensity could also be influenced by norepinephrine (Patriarchi et al., 2018), some of the lifetime signals could also be norepinephrine binding. We chose doublet lenses here because the 2.6x magnification provided more pixels per cell during imaging, therefore increasing the accuracy of fluorescence lifetime estimates per cell. No guide cannulae were used for these experiments. A titanium head plate was centered over the GRIN lens and fixed to the skull using Metabond. A 3D-printed PLA funnel was cemented onto the headplate to enable light shielding during experiments. The area surrounding the lens was covered with Metabond and then dark dental cement to reduce autofluorescence. The GRIN lens was protected by a cut-off tip of an Eppendorf tube (Fisher) that was secured using Kwik-Cast (WPI).

For overexpression of Tyrosine Hydroxylase in AVPV/PVpo dopaminergic neurons, pAAV-EF1α-DIO-TH-p2A-mRFP was bilaterally injected into the AVPV of TH-Cre mice (150 nl; Bregma: AP 0.25 mm, ML 0.3 mm, DV −5.5 mm). For a control condition, AAV1-hSyn-DIO-mCherry (150 nl; Addgene) was bilaterally injected into the AVPV of TH-Cre mice (150 nl; Bregma: AP 0.25 mm, ML 0.3 mm, DV −5.5 mm).

For knock down expression of the vesicular GABA transporter VGAT, pAAV-hsyn-flex-dsRed-shvgat (150 nl; Children’s Hospital Vector Core and Addgene 67845) was bilaterally injected into the AVPV of TH-Cre mice (150 nl; Bregma: AP 0.25 mm, ML 0.3 mm, DV −5.5 mm). For a control condition, AAV2/8 hSyn-flex-dsRed-shscramble (150 nl; Children’s Hospital Vector Core and Addgene 71383) was bilaterally injected into the AVPV of TH-Cre mice (150 nl; Bregma: AP 0.25 mm, ML 0.3 mm, DV −5.5 mm).

### Head-fixed photometry

Head-fixed photometry experiments were conducted as described previously (Zhang et al., 2021, 2024) using mice running on a circular treadmill (Burgess et al., 2016) in order to reduce potential motion artifacts. All photometry and optogenetic excitation lights were modulated as interleaved pulses controlled by the Nanosec photometry-behavioral system that was initially developed in a previous study (Zhang et al., 2021, 2024) (https://github.com/xzhang03/NidaqGUI). Each cycle (20 ms per cycle, or 50 cycles per second) consists of two steps. Step 1 is turning on the photometry (465 nm) LED for 6 ms and Step 2 is turning off the photometry LED for 14 ms. If Chrimson stimulation was being used, once every 5 cycles, the 630-nm LED was turned on immediately after Step 2 and then turned off 10 ms later, which corresponds to 10 Hz, 10 ms pulses (duty cycle = 10%). Recordings were conducted in complete darkness and a black heat shrink was placed around the fiber to circumvent the LED light from being seen by the mice. Each mouse was habituated to handling and head-fixation for at least two weeks before photometry experiments.

For experiments using a green sensor (e.g., dLight 1.1, iGABASnFR), excitation light from a 465-nm LED (for fluorescent sensor excitation, ∼100 μW peak) and a 630-nm LED (for optogenetic stimulation of Chrimson; 1 mW peak if used) were combined in a four-port fluorescent mini-cube (FMC4_E(460-490)_F(500-550)_O(580-650)_S, Doric) and transmitted to the implanted fiber via a patch cord (1 m length, NA 0.57, Doric). The emitted light was measured from the emission port of the mini-cube using a femtowatt photoreceiver (2151, Newport).

### Hormonally priming female mice

To increase female receptivity and prevent pregnancy from mating experiments, female mice were hormonally primed before being introduced to males in the experiments. Using a standard protocol (Mosig & Dewsbury, 1976; Wu et al., 2009), we subcutaneously injected WT females with 18-35 μg of 17β-estradiol benzoate (dissolved in sterile sesame oil, final volume: 50-100 μl) 48 hours before introduction of the female to the male, and injected them again with 50-100 μg progesterone (dissolved in sterile sesame oil, final volume 50-100 μl) 5-12 hours before introduction of the female to the male. The efficacy of the hormonal priming was spot-checked using the criteria in Byers et al. (Byers et al., 2012).

### Home-cage mating experiments

All mating behavior experiments were conducted during the circadian dark hours inside the home cages of the males. Food bowls, huts, running wheels, and any other materials that could obstruct the view of the inside of the cage were temporarily removed for the duration of the experiments. The cages were then placed in a dark environment illuminated only with IR lamps. Videos of the cage were taken with Flea3 cameras and Flycapture 2 (FLIR systems) at 30 Hz. Hormonally primed females were gently lowered into the cage in a transparent cup (∼10 cm in diameter) under dim red light, which was subsequently turned off.

To measure a male’s mating behaviors, we used a 15-min assay that starts at female entry. To measure mating behaviors without satiating the male’s mating drive (i.e., transfer of reproductive fluids), we typically either stopped the recording and separated the animals after the male gained intromission (i.e. last step in the sequence of mating behaviors before transfer of fluids), or gently shook the cage to separate them in the cases for which we could not stop the video. To satiate a male’s mating drive, hormonally primed females were placed in the home cage of the male mouse for 24 hours, and were spot-checked for mating plugs afterward. In experiments where we measured a male’s mating behaviors throughout recovery from satiety, we ensured that the male was never paired with the same female more than once.

### LED control for home cage optogenetic experiments

Light from the 465-nm LED was transmitted to implanted optic fibers via a fiber optic patch cord (400 μm diameter core, multimode, 2 m length, NA 0.57, Doric) that was secured to the fiber using zirconia sleeves (Precision Fiber Products). Black heat shrink was placed around the fiber to attempt to decrease the LED light seen by the mice. For optogenetic axon and soma stimulation, the LED pulses were turned on 1 min before female entry and turned off 5 min after female entry. LED control for optogenetic experiments was done using the Nanosec photometry system as above (https://github.com/xzhang03/NidaqGUI).

### Two-photon imaging

Two-photon imaging was performed using a two-photon resonant-galvo scanning microscope (NeuroLabWare) controlled by Scanbox (https://scanbox.org/) as described previously (Burgess et al., 2016; Zhang et al., 2021, 2024). An InSight X3 laser (Spectr-Physics) was used to excite the fluorophores (910-1050 nm), and the emission light was filtered (Green: 510/84 nm; Red: 607/70 nm; Semrock) before collection with the photomultiplier tubes (PMT) (H10770B-40; Hamamatsu). The XY scanning was performed using resonant/galvo mirrors and the Z scanning was achieved with an electrically tunable lens (Optotune).

Two-photon Fluorescence-lifetime Imaging Microscopy (FLIM) was enabled by modifying the existing two-photon microscope (NeuroLabWare). Both FLIM and the conventional two-photon imaging used the same excitation light path. In experiments involving FLIM, an extra beam-splitter (75T/25R, Semrock) was added to the emission light path such that 75% of the emission light was directed to the standard intensity-based PMT (H10770B-40; Hamamatsu) and the rest (25%) was split to the FLIM light path. In the FLIM light path, the emission light was filtered (510/84 nm; Semrock) before entering the hybrid PMT (Becker and Hickl). PMT data were digitized using a timecorrelated single-photon counting board (Becker and Hickl) to estimate photon counts and arrival times. Frames were constructed from the pixel and frame clocks in the NeuroLabWare microscope. Laser clocks were recorded with an independent photodiode (Becker and Hickl) in the excitation path. In the FLIM experiments, the laser intensity was controlled (<10 mW) so that the FLIM PMT received 2.5-5 × 105 photons per second (to avoid cross-pulse photon contamination, see below). Each frame was collected from 2-5 seconds of continuous scanning so that it contained at least 1 million photons/mm2 to ensure accurate lifetime estimates while minimizing laser power.

### In vivo two-photon imaging experiments

In vivo two-photon imaging of the MPOA via a GRIN lens in head-fixed males was generally performed as described previously (Lutas et al., 2019; Zhang et al., 2021, Zhang et al., 2024). The excitation wavelength was 910-920 nm. The experiments were carried out using head-fixed mice that could run freely on a circular treadmill (Burgess et al., 2016; Zhang et al., 2021, Zhang et al., 2024). Imaging was performed with a 4x 0.2 NA air objective (Nikon) in mice implanted with a doublet GRIN lens (NEM-050-25-10-860-DM, Grintech; NA 0.19 on the air side). Light shield material was used to protect the lens and the objective from external light. Imaging fields-of-view were at depths of 100-300 μm below the face of the GRIN lens. For each recording, we collected five 5-s FLIM frames of the same field of view to be averaged (10 seconds between frames).

### Electrophysiology in acute slices

To prepare ex vivo brain slices, 6-10 week-old TH-Cre mice that had been injected bilaterally in AVPV with a Cre-dependent mCherry were deeply anesthetized with isoflurane before decapitation and removal of the entire brain. Brains were immediately submerged in ice-cold, carbogen-saturated (95% O2, 5% CO2) choline-based cutting solution consisting of (in mM): 92 choline chloride, 10 HEPES, 2.5 KCl, 1.25 NaH2PO4, 30 NaHCO3, 25 glucose, 10 MgSO4, 0.5 CaCl2, 2 thiourea, 5 sodium ascorbate, 3 sodium pyruvate, oxygenated with 95% O2/5% CO2, measured osmolarity 310 – 320 mOsm/L, pH= 7.4. Then, 275-300 μm-thick coronal sections were cut with a vibratome (Campden 7000smz2) and incubated in oxygenated cutting solution at 34°C for 10 min. Next, slices were transferred to oxygenated ACSF (126 mM NaCl, 21.4 mM NaHCO3, 2.5 mM KCl, 1.2 mM NaH2PO4, 1.2 mM MgCl2, 2.4 mM CaCl2, 10 mM glucose) at 34°C for an additional 15 min. Slices were then kept at room temperature (20–24°C) for 45 min until use. A single slice was placed in the recording chamber where it was continuously superfused at a rate of 3–4 mL per min with oxygenated ACSF. Neurons were visualized with an upright microscope (SliceScope Pro 1000, Scientifica) equipped with infrared-differential interference contrast and fluorescence optics. Cell-attached recordings (seal resistance 20–50 MΩ) were made in voltage-clamp mode with the recording pipette filled with ACSF and holding current maintained at Vh = 0 mV.

All recordings were made using a Multiclamp 700B amplifier, and data were filtered at 2 kHz and digitized at 20 Hz. Access resistance (<30 MΩ) was continuously monitored by a voltage step and recordings were accepted for analysis if changes were <15%.

### Histology

Perfusion and histology were performed as described in Garfield et al (Garfield et al., 2016). Brain slices (60 μm thick) were collected and one of every three consecutive slices was scanned. The primary antibodies used in this paper are chicken anti-GFP (1:1000, Invitrogen, A10262, Lot 2035172 and 2089131; used for GFP), rat anti-mCherry (1:1000, ThermoFisher, M11217, Lot UJ287711; used for mCherry), rabbit anti-TH (1:500, Millipore, AB152, Lot 3443922; used for TH), and rabbit anti-VGAT (1:1000, Invitrogen, VGAT Recombinant Rabbit Monoclonal Antibody (HL1616), Lot ZI4476877; used for VGAT-mRNAi). The secondary antibodies used in this paper are donkey anti-rabbit 594 (1:1000, Invitrogen, A21207, Lot 21450222), donkey anti-chicken 488 (1:1000, Jackson, 703-545-155, Lot 138498), and donkey anti-rat 594 (1:1000, Jackson, 712-585-153, Lot 134903).)

### iDISCO

Clearing of the hypothalamus was generally performed as described in Renier et al. on a ∼2×2×4 to ∼2×2×8 mm3 chunk of the mouse hypothalamus (Renier et al., 2014). After clearing, the hypothalamus was mounted in a 3D-printed chamber as suggested in the iDISCO protocol. Using the iDISCO protocol allowed for imaging of the entire hypothalamus, ensuring that the small AVPV/PVpo nuclei were fully preserved and avoiding potential cell loss that may occur if sectioned. Tyrosine hydroxylase expression was identified with the antibodies: rabbit anti-TH (Millipore, AB152) and donkey anti-rabbit 594 (Invitrogen, A21207). In brief, samples of mouse hypothalamus undergo dehydration with a methanol series (20%, 40%, 60%, 80%, 100%), followed by an overnight incubation in 66% dichloromethane (DCM)/ 33% Methanol. Samples were then bleached overnight in 5% H2O2 in methanol at 4°C, and then rehydrated to with a methanol series to PBS (80%, 60%, 40%, 20%, PBS). Immunolabeling was performed by incubating samples ∼36 hours in permeabilization (11.5g of Glycine, 100mL of DMSO, 400mL PTx.2 [100mL 10X PBS, 2mL TritonX-100]) then blocking solution (42mL PTx.2, 3mL of Donkey Serum, 5mL of DMSO) at 37°C, followed by staining with primary and secondary antibodies in PTwH buffer (100mL 10X PBS, 2mL Tween-20, 1mL of 10mg/mL Heparin stock solution). Immunolabeled tissue was then dehydrated in methanol series (20%, 40%, 60%, 80%, 100%), incubation DCM, and immersed in dibenzyl ether for optical clearing before imaging. A 3D volume scan of TH expression was obtained using two-photon microscopy using a combination of scanning with resonant mirror (X), galvo mirror (Y), and electrically tunable lens (Z, axial).

### Quantification of sperm counts

After mice were perfused with PBS, the peritoneum was exposed until the testicular fat pads were visible. Bilateral cauda epididymis and the connected vas deferens were located by gently pulling on the testicular fat pads and dissected out. Each cauda epididymis was placed in a well with 3 mL PBS, sliced in half, and further diced into fine pieces. The suspension was mixed thoroughly with a pipette, and 1 mL of the suspension was transferred to an Eppendorf tube. A cell staining kit (Fisher, L7011) was used to stain and visualize the sperm cells. 3 μL bovine serum albumin, 5 μL propidium iodide, and 1 μL SPYR14 were added to the tube and mixed, followed by incubation in a 37°C water bath for 5–10 minutes. 3-5 μL of the samples were placed onto a microscope slide, covered with a coverslip, and imaged under a slider scanner.

### Data Analysis

All data analyses were performed using custom scripts in MATLAB (MathWorks), Python, ImageJ (NIH), SPCImage (Becker and Hickl), and Prism (GraphPad).

### Analyses of male mating behaviors

The onset of a bout of sniffing was defined as the moment when a male initiated a bout of movement toward the female that culminated in the male beginning to sniff the female (usually either around the face or anus). Since the individual sniffs were fast, we inferred that the sniffing bout was terminated when the male made one of the following actions: turning his head away from the female, turning his body away from the female, moving forward but in a different direction from the female, remaining still when the female moved forward, or beginning other non-sniffing behaviors. The onset of a bout of mounting was defined as the moment when the male climbed over a female and began rapid, shallow lower body thrusting, and terminated when the male was no longer over the female. During a bout of mounting, the male could gain intromission, which was evident from the male’s thrusting movements becoming deeper and slower (usually about once per 1-2 sec). The transfer of fluids (i.e., ejaculation) was marked by a specific series of events where an intromitted male began to shake for a few seconds, then fell on his side, then became temporarily immobilized even when the female moved away, and then stood up again to clean himself. When we spot-checked mating plugs after the assays, the results consistently matched the behaviors of the male.

To measure a male’s appetitive mating behavior, which is commonly defined as the sniffing of females (Ball & Balthazart, 2008), we calculated the fraction of time that a male spent in sniffing females in the time period before the first mount, or up to five minutes from the moment that the female is introduced, whichever came first. We used this 5-minute maximum time window because most sniffing occurred in the minutes immediately after female entry and, by focusing on the pre-mounting time, we avoided the situation where a male who began mounting early (within the 5-minute sniffing time window) or spent more time mounting may paradoxically spend less time sniffing. We chose five minutes as the maximum time window, because it roughly matched the mean latency to the first mounts by wild-type males (5.46 min). We note that, on average, animals with a fiber in the MPOA show ∼10% lower values in both the appetitive sniffing metric and the consummatory mounting metric than animals without a fiber in the MPOA, possibly due to minor damage to this region. Note, however, that the experiments involving optic fibers mostly involved within-animal comparisons.

To measure a male’s consummatory mating behavior, we calculated the fraction of trials in which a male began mounting in the first 15 minutes after female entry.

### Analyses of head-fixed photometry experiments

Head-fixed photometry data were analyzed as described previously (Zhang et al., 2021, Zhang et al., 2024). Pulses that trigger the photometry LED (50 Hz, see above) were used to determine when the LED was on. For each pulse (6 ms), we took the median of the corresponding data points in the photodetector trace to quantify the photometry signal. This results in a 50 Hz trace which we then filtered with a 10 Hz low-pass filter. Link (https://github.com/xzhang03/Photometry_analysis)

To estimate bleaching, we fitted the pre-stimulation data (first 4 min of each experiment) with a mono-exponential function and subtracted the estimated values from each timepoint in the photometry trace across the entire recording. Since all photometry experiments have a trial structure, we used the baseline fluorescence (e.g., the 10 s pre-stimulation window) to estimate ΔF/F0.

### Pre-processing of two-photon FLIM data

Pre-processing of FLIM data was performed as previously described (Zhang et al., 2021, Zhang et al., 2024), using an open-source software package in MATLAB (https://github.com/xzhang03/SPC_analysis). Because dLight1.1 was expressed non-cell-type-specifically, which resulted in a diffuse expression pattern, we used the entire field of view (FOV) for each lifetime calculation instead of segmenting individual cells. Pixels corresponding to high-autofluorescence and low-lifetime (<1000 ps) particles (which we presume to be immune-related debris) are excluded from the calculation. Each histogram was constructed from the aggregate photon arrival times in each FOV, usually from 300,000 – 1,000,000 photons per FOV. For each histogram, more than 98.5% of the photons can be modeled using a bi-exponential fit, indicating low (∼1.5%) contribution from autofluorescence (e.g., Fig. S2A). The first-moment method (Tm) and the integral extraction method (IEM) were applied to estimate lifetimes (D.-U. Li et al., 2008; Yasuda, 2006; Zhang et al., 2021, 2024), using equations shown in Fig. S2A. In both cases, lifetimes were plotted as changes from the pre-satiety baseline.

### Electrophysiology analysis

Cell-attached firing-rate recordings were analyzed using MATLAB by finding local peaks with pre-determined thresholds on current, charge (time-domain integral of current), and widths (https://github.com/xzhang03/ephys_analysis). The results were spot-checked manually to ensure fidelity.

### 3D quantification of TH+ cells in AVPV/PVpo

Two-photon image stacks were flattened and locally normalized using MATLAB by computing local means and standard deviations within a defined neighborhood (https://github.com/xzhang03/Local-normalize). Segmentation of TH expression was performed using Cellpose2.0 (Pachitariu & Stringer, 2022) on the locally-normalized images. Examples can be seen in Video S1.

### Statistics and reproducibility

Two-tailed t-test and ANOVA were performed using Prism. For ANOVA, multiple comparisons were done using the Sidak posthoc correction. Also for ANOVA, F values (F) and degrees of freedom (DF) are indicated in Supplementary Table 1. For these tests, each mouse was treated as an independent sample. We typically averaged data from 2-3 tests per mouse, except for the behavioral experiments in which most animals were tested only once. For two-photon FLIM experiments, each field of view was treated as an independent sample (7 fields of view per mouse). Hypothesis testing of fractions was done with two-tailed Fisher’s exact tests in MATLAB, and multiple comparisons were adjusted using the Bonferroni correction. The fractions are based on the numbers of trials. We also note the numbers of animals in figure legends. For all tests, non-significant (ns) outcomes indicate that the null hypothesis cannot be rejected (i.e., p ≥ 0.05). Asterisks indicate *p<0.05, **p<0.01, ***p<0.001. Exact p-values are given whenever appropriate. For plots of fractions, error bars represent 95% Jeffrey’s confidence interval (95% c.i.). For all other plots, error bars represent the standard error of the mean (s.e.m.) or standard deviation (s.d.). For each panel where a representative image is shown, similar independent images from different mice were reproduced at least 3 times.

**Figure S1.**
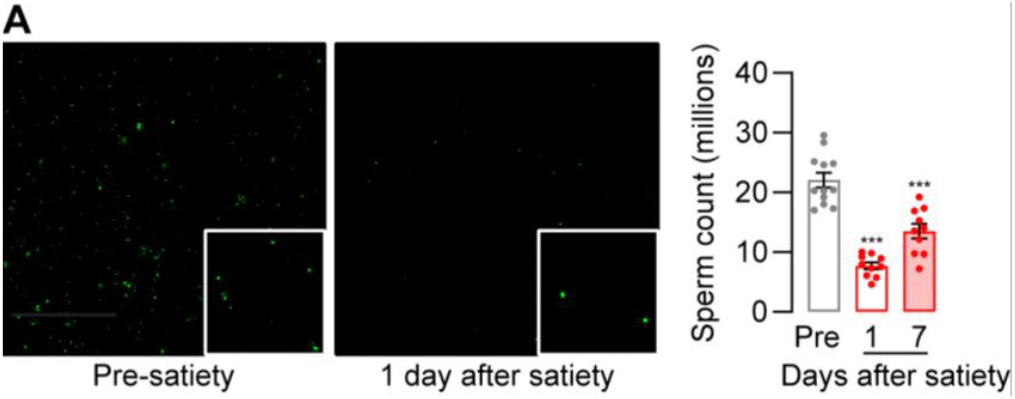
Recovery of mating behaviors tracks the recovery of sperm count. **(A)** After sexual satiation, sperm counts decrease and partially recover after 7 days. Sperm cells were extracted from dissected cauda epididymis and vas deferens, stained, and mounted on a microscope slide. Bottom right square shows a 5x zoomed in window. N = 12 male mice [Pre-satiety], 10 male mice [1 day after satiety], 10 male mice [7 days after satiety].

**Figure S2.**
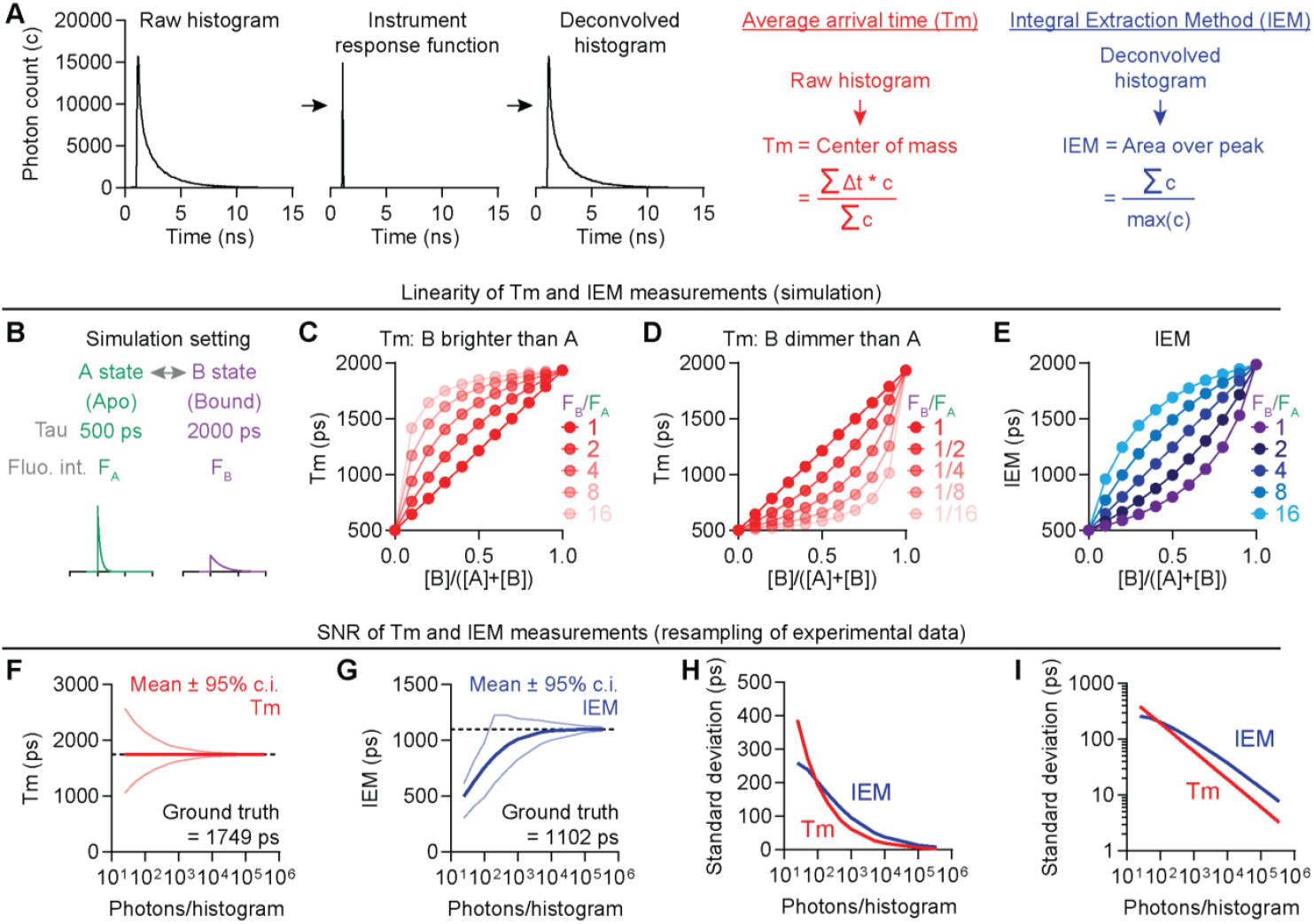
Two methods of fluorescent lifetime quantification. **(A)** Fluorescence lifetime quantification using average arrival time (Tm) and integral extraction method (IEM) metrics. In this example, the decay portion of the deconvolved histogram contains 335439 photons, 330850 (98.6%) of which can be captured with a double exponential fit, R^2^ = 0.998. **(B)** Setting for simulation: an idealized sensor has an apo, short-lifetime state (A state) and a ligand-bound, long-lifetime state (B state). Fluorescence intensity (Fluo. Int.) is denoted as F_A_ and F_B_. **(C)** Tm quantification of fluorescence lifetime only linearly reflects the absolute sensor activity (measured as the relative abundance of the bound state) if the long-lifetime state (B state) and the short-lifetime state (A state) are equally bright (i.e., F_B_/F_A_ = 1). If the long-lifetime state (B state) is brighter than the short-lifetime state (A state), the dose-response curve of Tm becomes left-shifted even if most sensor proteins are in the apo state. In this situation, the Tm metric tends to underestimate the dissociation constant (Kd) (i.e., overestimates sensor affinity). **(D)** If the long-lifetime state (B state) is dimmer than the short-lifetime state (A state), the dose-response curve of Tm becomes righted-shifted even if most sensor proteins are in the bound state. In this situation, Tm metric tends to overestimate Kd (i.e., underestimates sensor affinity). **(E)** IEM quantification of fluorescence lifetime only linearly reflects the absolute sensor activity (measured as the relative abundance of the bound state) if the brightness ratio of two states equals to the fluorescence-lifetime ratio (F_B_/F_A_ = Tau_B_/Tau_A_). If F_B_/F_A_ > Tau_B_/Tau_A_, IEM overestimates sensor affinity. If F_B_/F_A_ < Tau_B_/Tau_A_, IEM underestimates sensor affinity. **(F)** Increasing photon counts does not affect mean Tm values but it reduces noise. Data are generated by bootstrapping dLight1.1 fluorescence lifetime data for 10,000 iterations. In each iteration, the number of photons from the histogram (see A) is resampled, with replacement, to the desired counts (e.g., 1000 photons in histogram). Then, the corresponding lifetime metrices are calculated and plotted as mean ± 95% c.i. **(G)** At low photon counts, mean IEM values deviate significantly from ground truth. Increasing photon counts systematically change mean IEM values, make IEM estimates more accurate, and reduces noise. **(H-I)** Noise comparison using standard deviations of Tm and IEM estimates at different photon counts.

**Figure S3.**
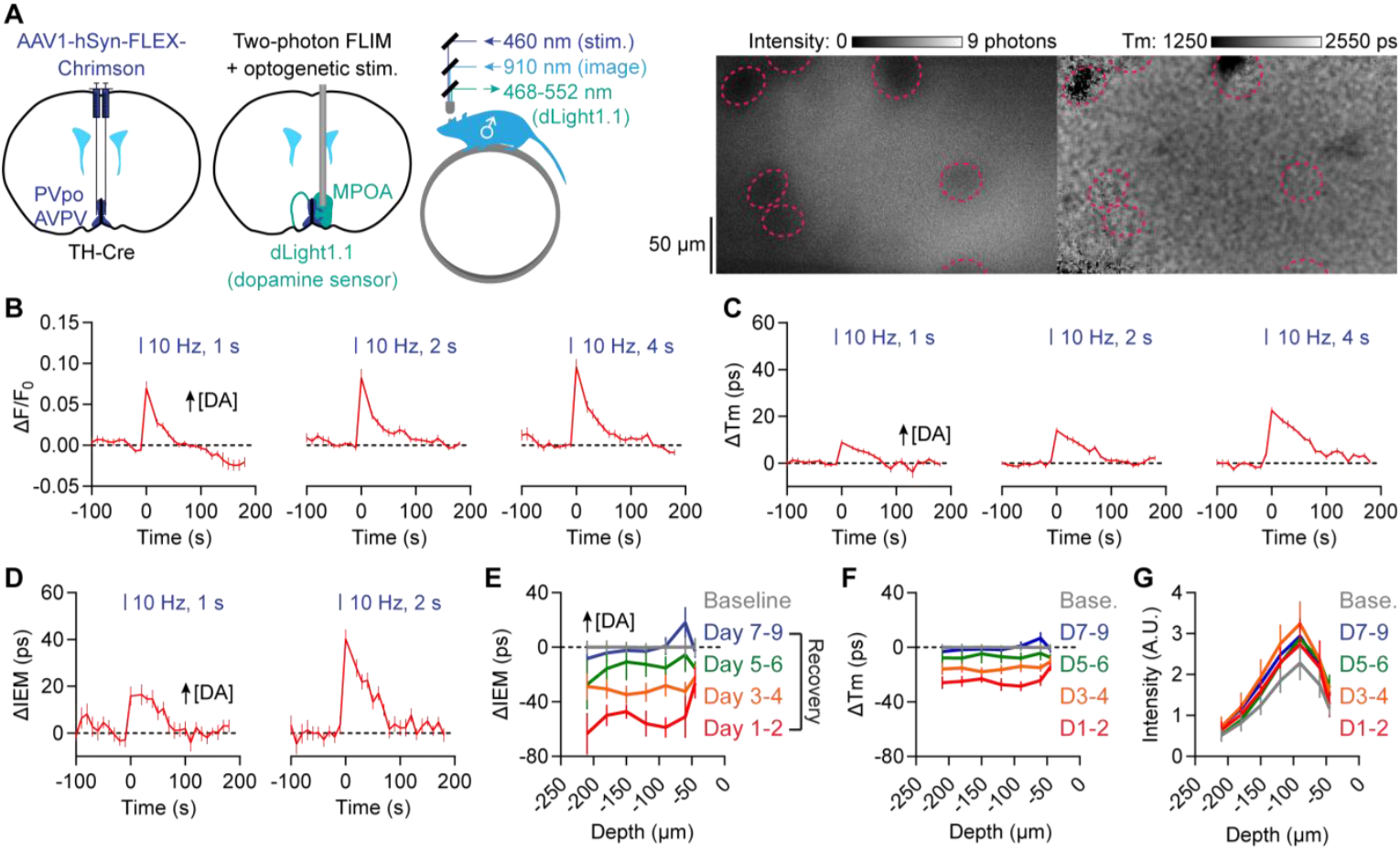
Experimental comparison of IEM and Tm lifetime quantifications. **(A)** Left: comparison of quantification methods using *in vivo* 2p-FLIM imaging of dopamine concentration (measured using dLight 1.1) in the MPOA during photostimulation of dopaminergic inputs from AVPV/PVpo neurons. Right: example fields of view of dLight1.1 intensity and Tm quantifications of fluorescence lifetime. Each Tm value was estimated using a local binning of 15×15 pixels. Dotted circles show dark spots of likely blood vessel locations. **(B)** Optogenetic stimulation of dopaminergic AVPV/PVpo axons triggers local dopamine release in the MPOA, measured using ΔF/F_0_ of dLight1.1 fluorescence intensity. Note the small but consistent effect of photobleaching on baseline dLight1.1 fluorescence intensity (also confirmed in longer unaveraged traces, not shown). N = 3 male mice. **(C-D)** IEM quantification (D), in comparison to Tm quantification (C), yields greater dLight 1.1 lifetime changes in response to optogenetic stimulation of AVPV/PVpo dopamine axons. N = 3 male mice. **(E-G)** Both IEM (E) and Tm (F) quantification of fluorescence lifetime reveal a significant decrease in tonic dopamine concentration after sexual satiation that gradually recovers to baseline across a week. However, this fall and recovery were not observed when using sensor intensity to quantify changes (G). N = 3 male mice, mean ± s.e.m.

**Figure S4.**
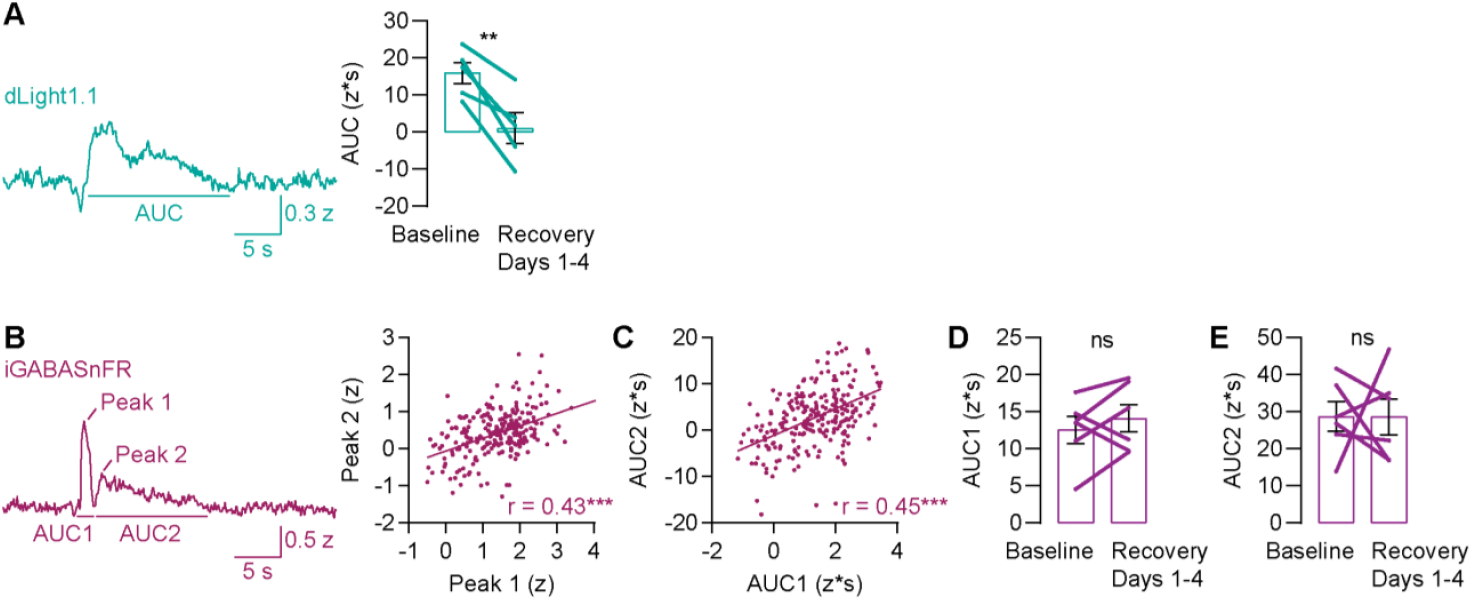
iGABASnFr photometry demonstrates biphasic evoked GABA release in the MPOA. **(A)** Sexual satiation reduced the size of the dlight1.1 transient evoked by photostimulating dopamine AVPV/PVpo axons in the MPOA in vivo. Transient sizes are quantified using areas under curve (AUC). N = 5 male mice. **(B-C)** Brief photostimulation of dopaminergic AVPV/PVpo axons in the MPOA *in vivo* (Figure 3F-3G) evoked a biphasic GABA response, measured by iGABASnFR photometry. The two peaks correlate on a trial-by-trial basis at the level of both peak ΔF/F_0_ (B) and AUC (C). The peaks were chosen on fixed windows of 0.3-2 seconds (Peak 1 and AUC1) and 2-14 seconds (Peak 2 and AUC2) after stimulation offset. A 10-hz lowpass filter was used to smooth individual traces before the calculations. N = 244 trials form 6 male mice. **(D-E)** Neither of the two iGABASnFR peak sizes changed significantly after sexual satiation. N = 6 male mice. Mean ± s.e.m. *p<0.05, **p<0.01, ***p<0.001. See Supplementary Table 1 for statistics.

**Figure S5.**
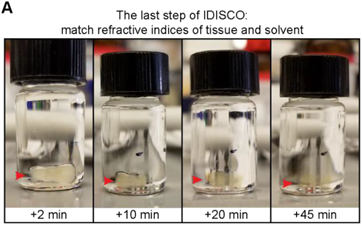
Clearing of mouse hypothalamus using iDISCO. **(A)** A ∼2 × 2 × 8 mm^3^ chunk of the mouse hypothalamus (red arrow head) gradually became clear during the last step of iDISCO. During this step, an organic solvent (dibenzyl ether) was used to match the refractive index of the preprocessed hypothalamus tissue, thus reducing light scattering and making the tissue transparent. After initial testing, the chunk size was shrunk down to 2 × 2 × 2 mm^3^ to reduce clearing time.

