## Supplementary Table 1 for "Slow-Timescale Regulation of Dopamine Release and Mating Drive Over Days"

**Supplementary Table 1. Statistical analysis**

| Figure | N_total | P values | F and DF | Statistical notes |
|---|---|---|---|---|
| 1B | 11 males (Pre)<br>11 males (0 days)<br>11 males (1 day)<br>11 males (3 days)<br>11 males (5 days)<br>11 males (7 days)<br>11 males (9 days) | p < 0.0001 (0 days)<br>p < 0.0001 (1 days)<br>p < 0.0001 (3 days)<br>p = 0.431 (5 days)<br>p = 0.993 (7 days)<br>p = 0.997 (9 days) | F = 134.8<br>DF = 6 | Two-tailed one-way ANOVA with Sidak post hoc test.<br>Comparisons are with the pre-satiety group. |
| 1C | 33 trials (Pre)<br>11 trials (0 days)<br>22 trials (1 day)<br>22 trials (3 days)<br>22 trials (5 days)<br>22 trials (7 days)<br>22 trials (9 days) | p < 0.0001 (0 days)<br>p < 0.0001 (1 days)<br>p < 0.0001 (3 days)<br>p = 0.032 (5 days)<br>p = 0.981 (7 days)<br>p = 0.933 (9 days) | DF = 6 | Two-tailed Fisher's exact test with Bonferroni correction.<br>Comparisons are with the pre-satiety group. |
| 1D | 11 males (Pre)<br>11 males (0 days)<br>11 males (1 day)<br>11 males (3 days)<br>11 males (5 days)<br>11 males (7 days)<br>11 males (9 days) | p = 0.998 (0 days)<br>p = 0.731 (1 days)<br>p = 0.973 (3 days)<br>p = 0.315 (5 days)<br>p = 0.232 (7 days)<br>p = 0.857 (9 days) | F = 3.1<br>DF = 6 | Two-tailed one-way ANOVA with Sidak post hoc test.<br>Comparisons are with the pre-satiety group. |
| 1F | 9 FOVs from 3 males | N/A | N/A | N/A |
| 1G | 6 FOVs from 3 males (pre)<br>6 FOVs from 3 males (1 day)<br>6 FOVs from 3 males (3 days)<br>6 FOVs from 3 males (5 days)<br>6 FOVs from 3 males (7 days) | p < 0.0001 (1 days)<br>p < 0.0057 (3 days)<br>p = 0.320 (5 days)<br>p = 0.999 (7 days) | F = 13.4<br>DF = 4 | Two-tailed one-way ANOVA with Sidak post hoc test.<br>Comparisons are with the pre-satiety group. |
| 2A | Example neuron | N/A | N/A | N/A |
| 2B | Example neuron | N/A | N/A | N/A |
| 2C | Example neuron | N/A | N/A | N/A |
| 2D | 43 neurons from 7 males | N/A | N/A | N/A |
| 2E | 58 neurons from 8 males | N/A | N/A | N/A |
| 2F | 48 neurons from 7 males | N/A | N/A | N/A |
| 2G | 28 neurons from 7 males (baseline)<br>36 neurons from 8 males (1 day)<br>15 neurons from 7 males (2 days) | p = 0.008 (baseline vs. 1 day)<br>p = 0.270 (baseline vs. 2 days)<br>p = 0.0002 (1 day vs. 2 days) | F = 10.1<br>DF = 2 | Two-tailed one-way ANOVA with Sidak post hoc test. |
| 2H | 15 neurons from 7 males (baseline)<br>22 neurons from 8 males (1 day)<br>33 neurons from 7 males (2 days) | p = 0.010 (baseline vs. 1 day)<br>p = 0.425 (baseline vs. 2 days)<br>p = 0.006 (1 day vs. 2 days) | F = 8.2<br>DF = 2 | Two-tailed one-way ANOVA with Sidak post hoc test. |
| 3A left | 7 males (no stim.)<br>9 males (soma stim.)<br>9 males (axon stim.) | p = 0.003 (no stim. vs. soma stim.)<br>p = 0.036 (no stim. vs. axon stim.)<br>p = 0.389 (soma stim. vs axon stim.) | F = 7.7<br>DF = 2 | Two-tailed one-way ANOVA with Sidak post hoc test. |
| 3A right | 7 trials (no stim.)<br>9 trials (soma stim.)<br>9 trials (axon stim.) | p = 0.005 (no stim. vs. soma stim.)<br>p = 0.009 (no stim. vs. axon stim.)<br>p = 0.907 (soma stim. vs axon stim.) | DF = 2 | Two-tailed Fisher's exact test with Bonferroni correction. |
| 3B left | 7 males (no stim.)<br>9 males (soma stim.)<br>9 males (axon stim.) | p = 0.690 (no stim. vs. soma stim.)<br>p = 0.110 (no stim. vs. axon stim.)<br>p = 0.404 (soma stim. vs axon stim.) | F = 2.3<br>DF = 2 | Two-tailed one-way ANOVA with Sidak post hoc test. |
| 3B right | 14 trials (no stim.)<br>18 trials (soma stim.)<br>18 trials (axon stim.) | p = 0.889 (no stim. vs. soma stim.)<br>p = 0.951 (no stim. vs. axon stim.)<br>p = 0.999 (soma stim. vs axon stim.) | DF = 2 | Two-tailed Fisher's exact test with Bonferroni correction. |
| 3C left | 7 males (no stim.)<br>9 males (soma stim.)<br>9 males (axon stim.) | p = 0.951 (no stim. vs. soma stim.)<br>p = 0.191 (no stim. vs. axon stim.)<br>p = 0.288 (soma stim. vs axon stim.) | F = 1.9<br>DF = 2 | Two-tailed one-way ANOVA with Sidak post hoc test. |
| 3C right | 14 trials (no stim.)<br>18 trials (soma stim.)<br>18 trials (axon stim.) | p = 0.771 (no stim. vs. soma stim.)<br>p = 0.829 (no stim. vs. axon stim.)<br>p = 0.364 (soma stim. vs axon stim.) | DF = 2 | Two-tailed Fisher's exact test with Bonferroni correction. |
| 3E | 5 males (baseline)<br>5 males (1-4 days) | N/A | N/A | N/A |
| 3F | 6 males (baseline)<br>6 males (1-4 days) | N/A | N/A | N/A |
| 4B | 10 males (baseline)<br>9 (1-3 days)<br>7 (4-6 days)<br>7 (7-9 days) | N/A | N/A | N/A |
| 4C | 9 males (Pre, TH-OE)<br>9 males (0 days, TH-OE)<br>9 males (1 day, TH-OE)<br>9 males (3 days, TH-OE)<br>8 males (Pre, mCherry)<br>8 males (0 days, mCherry)<br>8 males (1 day, mCherry)<br>8 males (3 days, mCherry) | p < 0.0001 (0 days, TH-OE)<br>p < 0.0001 (1 day, TH-OE)<br>p = 0.358 (3 days, TH-OE)<br>p < 0.0001 (0 days, mCherry)<br>p < 0.0001 (1 day, mCherry)<br>p < 0.0001 (3 days, mCherry) | F = 35.9<br>DF = 7 | Two-tailed one-way ANOVA with Sidak post hoc test.<br>Comparisons are with the pre-satiety group. |

| Figure | N_total | P values | F and DF | Statistical notes |
|---|---|---|---|---|
| 4D | 27 trials (Pre, TH-OE)<br>9 trials (0 days, TH-OE)<br>18 trials (1 day, TH-OE)<br>18 trials (3 days, TH-OE)<br>24 trials (Pre, mCherry)<br>8 trials (0 days, mCherry)<br>8 trials (1 day, mCherry)<br>16 trials (3 days, mCherry) | p < 0.0001 (0 days, TH-OE)<br>p = 0.221 (1 day, TH-OE)<br>p = 0.74 (3 days, TH-OE)<br>p < 0.0001 (0 days, mCherry)<br>p < 0.0001 (1 day, mCherry)<br>p < 0.021 (3 days, mCherry) | DF = 7 | Two-tailed Fisher’s exact test with Bonferroni correction.<br>Comparisons are with the pre-satiety group. |
| 4E | 8 males (Pre, VGAT-RNAi)<br>8 males (0 days, VGAT-RNAi)<br>8 males (1 day, VGAT-RNAi)<br>8 males (3 days, VGAT-RNAi)<br>8 males (Pre, scramble)<br>8 males (0 days, scramble)<br>8 males (1 day, scramble)<br>8 males (3 days, scramble) | p < 0.0001 (0 days, VGAT-RNAi)<br>p = 0.0007 (1 day, VGAT-RNAi)<br>p = 0.939 (3 days, VGAT-RNAi)<br>p < 0.0001 (0 days, scramble)<br>p < 0.0001 (1 day, scramble)<br>p < 0.0001 (3 days, scramble) | F = 36.2<br>DF = 7 | Two-tailed one-way ANOVA with Sidak post hoc test.<br>Comparisons are with the pre-satiety group. |
| 4F | 24 trials (Pre, VGAT-RNAi)<br>8 trials (0 days, VGAT-RNAi)<br>16 trials (1 day, VGAT-RNAi)<br>16 trials (3 days, VGAT-RNAi)<br>24 trials (Pre, scramble)<br>8 trials (0 days, scramble)<br>16 trials (1 day, scramble)<br>16 trials (3 days, scramble) | p < 0.0001 (0 days, VGAT-RNAi)<br>p = 0.0002 (1 day, VGAT-RNAi)<br>p = 0.517 (3 days, VGAT-RNAi)<br>p < 0.0001 (0 days, scramble)<br>p < 0.0001 (1 day, scramble)<br>p < 0.0001 (3 days, scramble) | DF = 7 | Two-tailed Fisher’s exact test with Bonferroni correction.<br>Comparisons are with the pre-satiety group. |
| S1A | 12 males (Pre)<br>10 males (1 day)<br>10 males (7 days) | p < 0.0001 (1 day)<br>p < 0.0001 (7 days) | F = 46.2<br>DF = 2 | Two-tailed one-way ANOVA with Sidak post hoc test.<br>Comparisons are with the pre-satiety group. |
| S3B | 9 FOVs from 3 males | N/A | N/A | N/A |
| S3C | 9 FOVs from 3 males | N/A | N/A | N/A |
| S3D | 9 FOVs from 3 males | N/A | N/A | N/A |
| S3E | 6 FOVs from 3 males (pre)<br>6 FOVs from 3 males (1 day)<br>6 FOVs from 3 males (3 days)<br>6 FOVs from 3 males (5 days)<br>6 FOVs from 3 males (7 days) | N/A | N/A | N/A |
| S3F | 6 FOVs from 3 males (pre)<br>6 FOVs from 3 males (1 day)<br>6 FOVs from 3 males (3 days)<br>6 FOVs from 3 males (5 days)<br>6 FOVs from 3 males (7 days) | N/A | N/A | N/A |
| S3G | 6 FOVs from 3 males (pre)<br>6 FOVs from 3 males (1 day)<br>6 FOVs from 3 males (3 days)<br>6 FOVs from 3 males (5 days)<br>6 FOVs from 3 males (7 days) | N/A | N/A | N/A |
| S4A | 5 males | p = 0.008 | DF = 4 | Two-tailed t-test. |
| S4B | 244 trials | p < 0.0001 | N/A | P value for non-zero correlation coefficient |
| S4C | 244 trials | p < 0.0001 | N/A | P value for non-zero correlation coefficient |
| S4D | 6 males | p = 0.399 | DF = 5 | Two-tailed t-test. |
| S4E | 6 males | p = 0.992 | DF = 5 | Two-tailed t-test. |
